# A fast algorithm for 3D volume reconstruction from light field microscopy datasets

**DOI:** 10.1101/2023.03.23.533935

**Authors:** Jonathan M. Taylor

## Abstract

Light field microscopy can capture 3D volume datasets in a snapshot, making it a valuable tool for high-speed 3D imaging of dynamic biological events. However, subsequent computational reconstruction of the raw data into a human-interpretable 3D+time image is very time-consuming, limiting the technique’s utility as a routine imaging tool. Here we derive improved equations for 3D volume reconstruction from light field microscopy datasets, leading to dramatic speedups. We characterise our open-source Python implementation of these algorithms, and demonstrate real-world reconstruction speedups of more than an order of magnitude compared to established approaches. The scale of this performance improvement opens up new possibilities for studying large timelapse datasets in light field microscopy.

---

Light field microscopy provides snapshot 3D imaging of dynamic scenes *via* a lenslet array placed in the image plane of the microscope, which casts a multi-apertured intensity pattern onto a camera sensor. The mix of spatial and angular information about the target sample emission in this single raw 2D snapshot image allows 3D image reconstruction across an extended depth-of-field (Fig. 1a), but only after intensive computational processing (1). Solving this inverse problem is extremely computationally demanding, traditionally requiring the computation of thousands of Fourier transforms per iteration. This requirement for high levels of computational resource is particularly problematic given the attractiveness of light field imaging for 3D time series (2–5), where a large number of different timepoints must all be reconstructed. Here we will demonstrate how to mathematically simplify the light field reconstruction process, speeding up real-world computation times by more than an order of magnitude, while delivering identical output volume results.

**Fig. 1.**
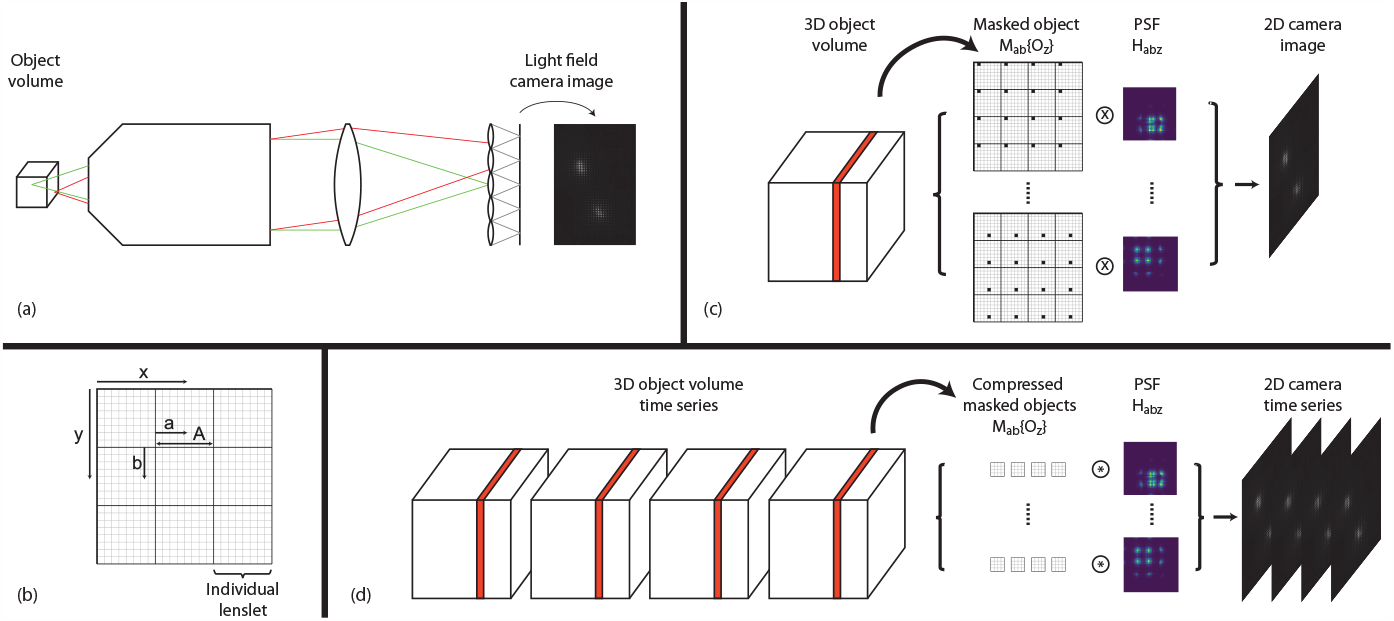
(a) Optical schematic of light field microscopy. (b) Pixel indexing relative to lenslet footprints. (c) Conventional projection operation for deconvolution (⊗ represents convolution). (d) New fast projection strategy (⊛ represents a new custom operation - see main text).

The image reconstruction process is an inverse problem that can be cast as a deconvolution. The spatially-variant point spread function (PSF) of the light field microscope is typically computed theoretically from wave-optics calculations (1, 2), and Richardson-Lucy deconvolution is then used to estimate the 3D volume that gives rise to the measured 2D intensity pattern observed on the camera sensor. The standard Richardson-Lucy algorithm is described by the following iterative formula:

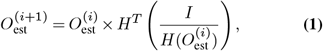

although the widely-used light field implementation in (2) uses an alternative variation (see footnote (6) for further discussion) where the error term is computed in object space:

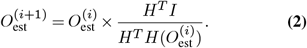

In either form, × denotes elementwise multiplication and the fraction implies elementwise division; 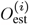 is the estimation of the 3D object to be reconstructed, at iteration *i*; *I* is the 2D light field image recorded on the camera; *H* is the “forward projection” operator mapping from the object *O* to the resultant camera image *I*; and *H*^*T*^ is the matrix transpose of the operator *H*. The optical interpretation of *H*^*T*^ leads to it being termed the backward-projection operator. A typical starting condition would be 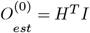,and *O*_est_ converges to an estimate of the true object *O* over *N*_iter_ ∼10 iterations.

The basic building blocks of the deconvolution problem are therefore the forward- and backward-projection operators, *H* and *H*^*T*^, which model the image formation process. In light field microscopy these projection operators are expressed as the sum of many separate convolution operations. Given a three-dimensional object *O* consisting of voxels indexed *O*_*xyz*_, each pixel value *I*_*mn*_ of a forward-projected image *I* can be computed as

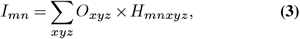

where *H*_*mnxyz*_ are the matrix elements of the PSF applicable to voxel *x, y, z*. To render the reconstruction problem tractable, raw images are resampled such that footprint of each lenslet in the lenslet array spans an exact odd integer number of camera pixels, *A*. Thus any subpixel *a, b* at the same relative position within any lenslet footprint will have the same the point spread function (Fig. 1b). This simplifies the problem, enabling Equation 3 to be rewritten as:

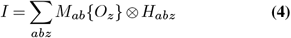

where *M*_*ab*_ is a masking operator which zeroes all pixels except those in the image pixel group satisfying *x* mod *A* = *a* and *y* mod *A* = *b*, and ⊗ is the convolution operator. *H*_*abz*_ represents the complete PSF for a voxel at coordinate *a, b, z*. Thus the forward-projected image from an object consisting of *Z* individual *z*-planes can be computed using a total of *A*^2^*Z* convolution operations.

According to the convolution theorem, each convolution consists of two Fourier transforms and one inverse, each computed using the Fast Fourier Transform (FFT) algorithm:

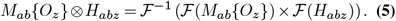

Any strategy aiming for more than merely incremental performance speedup of the overall calculation must speed up all three of the FFT operations in this equation. Speeding up any one of these on its own would be insufficient: even if the run time for one of them on its own could be reduced to zero, the overall run time would still only be improved by a factor of ∼30%. In what follows we will demonstrate how to improve the run time of each of these three operations in turn, to achieve an order of magnitude speedup in computation time. Our strategy is illustrated schematically in Fig. 1c-d.

The discrete Fourier transform of the masked object *M*_*ab*_{*O*_*z*_} involves a high degree of redundancy, since most elements of this masked object array are zero. We observe that this problem can be simplified by generalising the Danielson-Lanczos lemma (7, 8) to an *A*-way result. In one dimension, the *k*^th^ element *F*_*k*_ of the discrete Fourier transform of a function *f* can be expressed in terms of *A* smaller Fourier transforms:

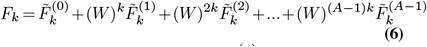

where *W* = exp(−2*πi/X*) and 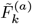 is the (reduced-size) discrete Fourier transform of 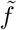, where 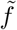 is a vector consisting of only the nonzero elements of *M*_*a*_{*f*}. Note that, in the notation of Equation 6, *k* indexes the reduced-size discrete Fourier transform 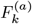 beyond its normal domain of *k* ∈ [0,*X*/*A*), exploiting the periodic boundary conditions implicit in the Fourier transform.

In our case the *M* operator ensures that only one of the *A* distinct terms on the right-hand size of Equation 6 is nonzero. Therefore, eliminating all the zero terms, generalising to two dimensions, and summing over all image pixel groups *a, b*:

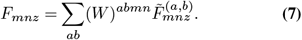

Consequently, instead of requiring *A*^2^ FFTs that each operate over the full image size *XY*, we have reduced these to operating on arrays of size *XY/A*^2^ (compressed arrays representing image pixel groups each containing only those pixels retained by the *M*_*ab*_{*O*_*z*_} operator). We have therefore reduced the computational requirements of these FFTs by a factor of *A*^2^. After computation of the FFTs, the weighting multiplications in Equation 7 must still be applied, albeit in an operation of reduced computational complexity 𝒪(*XY*), but overall the computational demands of computing *F*_*nm*_ are dramatically reduced.

We now move on to consider the Fourier transform of the point spread function *H*_*abz*_. *H* and its Fourier transforms are invariant properties of the imaging system. If sufficient memory storage is available, the Fourier transforms can be precomputed once and the computation of ℱ(*H*_*abz*_) eliminated completely from Equation 4. However, given typical values of *A* ≥15, *Z* ∼ 50 and megapixel images, tens or hundreds of GB of memory would be required to cache all the precomputed values. That may be feasible on some highend CPU-based platforms, but exceeds the capacity of most GPU platforms. Nevertheless, even when precalculation is not possible, our algorithm amortises the computation of each Fourier transform across batches of multiple timepoints in a time series dataset, reconstructing each batch concurrently. We also exploit symmetry relationships in the PSFs, to permit rapid computation of e.g. ℱ(*H*_(*A*− *a*)*bz*_) once ℱ(*H*_*abz*_) has been computed on-the-fly.

Finally, by explicitly substituting Equation 5 into Equation 4 and exploiting the distributive property of the Fourier transform, we can dramatically reduce the number of inverse Fourier transforms required:

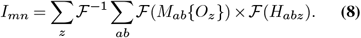

We note that in a practical implementation using a non-circulant Fourier transform it is not feasible to further promote the inverse FFT outside the summation over *z*, because the size of the intermediate arrays will vary according to the lateral extent of *H*_*abz*_, and hence vary with *z*.

The backward-projection operator required for Richardson-Lucy deconvolution is traditionally denoted *H*^*T*^ in recognition that it is the transpose of the matrix operator *H*. However, in practice *H* and *H*^*T*^ are both implemented as convolutions (as per Equations 3-4). Hence we follow an almost identical approach for the backward projection, starting from the following equation in place of Equation 4:

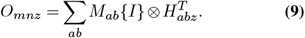

We note that during the one-time computation of *H* and *H*^*T*^ during initial optical modelling of the microscope PSF, computing *H*^*T*^ does not require additional convolution operations as used in (2), and can instead be populated near-instantaneously by pixelwise reshuffling of elements of *H*. To achieve our anticipated order-of-magnitude speedup by using Equation 7 to compute ℱ(*M*_*ab*_{*O*_*z*_}) in Equation 5, it was necessary to write carefully-optimized custom computer code, since the specific multiplication and tiling process underpinning a practical implementation of Equation 7 is a performance bottleneck, and it is a highly bespoke operation that is not available in standard numerical libraries. We have made available a high-performance open-source reference implementation of our complete light field reconstruction algorithm (9). It is written primarily in the Python programming language, with performance-critical code written in C++, cython (10) and CUDA, drawing on the FFTW library (11) to compute Fourier transforms in our C++ code. To ensure maximum performance our code incorporates elements of dynamic load-balancing between CPU cores, and machine-adaptive runtime optimisations. We also provide a demonstration of how it can be integrated as-is into existing Matlab workflows (12).

Table 1 presents performance benchmarks for our light field reconstruction code. For batch reconstruction of a light field timelapse series, our new approach delivers a performance gain of 8−35× compared to the widely-used MATLAB implementation (2) based on Equations 2, 4 and 9, and for maximum speed computation can be offloaded to a consumer GPU unit.

**Table 1.** Measured run times (in seconds per frame) for reconstructing a light field timelapse series, showing speedups of 8 − 35× compared to the existing state-of-the-art (Ref. (2)). The test parameters, representative of a typical light field imaging scenario, are specified in (9). Run times are also shown for a batch size of 1, as discussed in the main text, although that scenario does not allow our algorithms to perform at their fastest. System A: 8 core Intel Xeon, 3GHz, 32GB RAM. System A GPU: PNY Quadro RTX A4000, 1.56GHz, 16GB RAM, 224.03GB/s. System B: 16 core Intel Xeon, 3.2GHz, 256GB RAM. System C: 4 core Intel Core i5, 2.5GHz, 16GB RAM.

| System | CPU/<br>GPU | Batch<br>size | This work | (2) | Speedup |
| --- | --- | --- | --- | --- | --- |
| A | 8xCPU | 32 | **78.4** | 1075.9 | 14× |
| A | GPU | 24 | **5.6** | 194.5 | 35× |
| B | 16xCPU | 32 | **30.34** | - |  |
| C | 4xCPU | 32 | **140.0** | 1095.6 | 7.8× |
| A | 8xCPU | 1 | 628.8 | 1075.9 | (1.7×) |
| A | GPU | 1 | 19.0 | 194.5 | (10×) |

As explained above, our implementation performs optimally when batch-reconstructing multiple timepoints in a time series dataset simultaneously. Nevertheless, even with a non-optimal batch size of 1 we note that our implementation already outperforms existing implementations (Table 1). Fig. 2 explores the relationship between overall run time and batch size for our code, revealing a clear linear-plus-baseline scaling law. Regardless of batch size, a fixed amount of computational work must be performed to compute ℱ(*H*). This baseline work can only be eliminated completely if sufficient RAM is available to cache pre-calculated values for all ℱ(*H*), which can extend to hundreds of GB. The total additional work (computing the other elements of Equation 4 via Equations 7 and 8) scales linearly with the batch size. Hence there is a clear advantage to our amortising the constant work across simultaneous deconvolutions of multiple timepoints. In our GPU implementation we measure that the linear-scaling work is more substantial compared to the baseline work of computing ℱ(*H*), probably due to the challenges of developing optimized CUDA kernels to implement the highly specific custom operations embodied in Equation 7. This means that there are diminishing additional benefits to increasing the batch size beyond 8 on a GPU, which happily in turn means the RAM requirements are lower on a GPU, where RAM is typically scarcer.

**Fig. 2.**
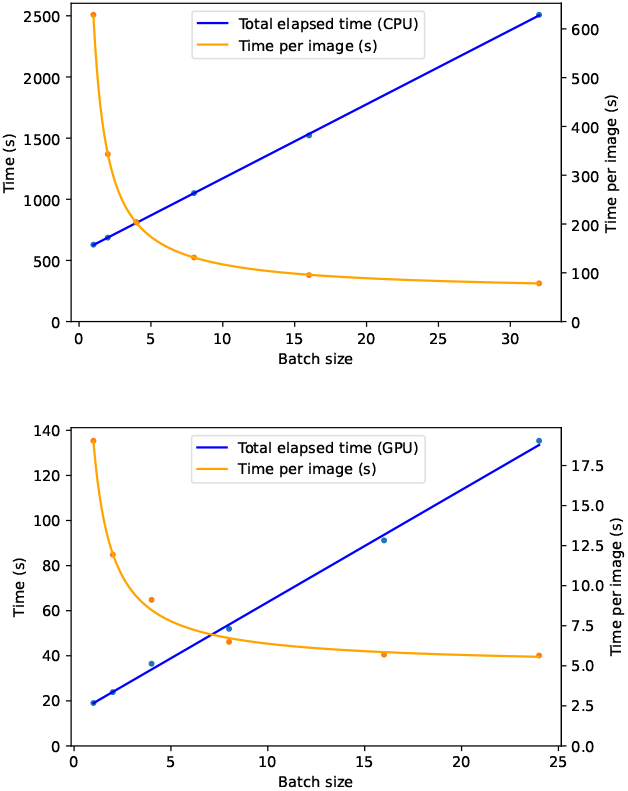
Performance scaling with batch size, showing trend of constant baseline work (computing ℱ(*H*)) plus linear scaling of additional work per batch item. Datapoints are measured run times (System A as specified in Table 1) and trendlines are a fit to a linear scaling model.

Fig. 3 confirms that our implementation scales well when the work is parallelised across multiple CPU cores: the benchmark run time with 16 parallel threads was over 12 times faster than the single-threaded run time. The small decrease in efficiency (i.e. speedup being slightly less than *n* times when using *n* threads) can be explained by the increased pressure that multithreaded code imposes on the system’s memory bandwidth. The slight anomaly in efficiency for 2 threads is likely related to the dual-CPU architecture of the testbed system.

**Fig. 3.**
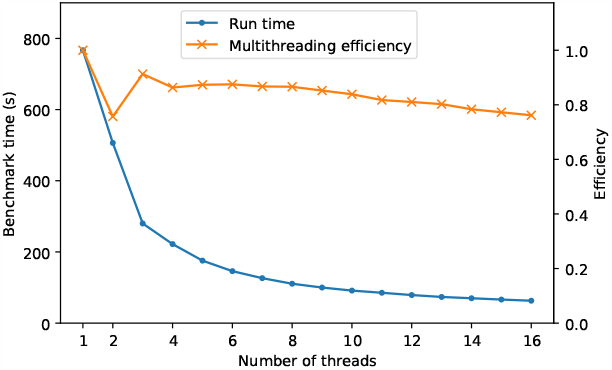
Performance scaling with number of parallel CPU threads, showing run-time improvement with increasing number of parallel threads (System B as specified in Table 1). Multithreading efficiency is defined as 1.0 for a case where the measured time for an 𝓃-thread scenario is 𝓃 times as fast as the 1-thread time.

While many researchers continue to prefer the mathematically precise reconstruction afforded by the classical approach of Equation 2, others are researching the use of machine-learning to estimate the object from the raw camera image (5, 13–15). The aim of such research is to compute an object estimate that is as faithful as possible to the true object, while using less computational time than required to explicitly compute Equation 2. Some approaches commence with a limited number of Richardson-Lucy iterations (13) or use direct optical modelling during the training phase (5); computational performance in both these scenarios will benefit directly from our new results. Pure machine-learning based approaches also stand to benefit from our results: machine-learning architectures in imaging are commonly based around convolutional neural networks, and it is increasingly recognised that networks perform better and can be trained faster when the network structure encapsulates physical insights into the image formation process (14, 15). Thus the mathematical insights behind our results hold promise for improving performance and effectiveness of future machinelearning architectures for light field microscopy reconstruction.

In summary, we have presented mathematical results enabling a dramatic reduction in computational work for 3D image reconstruction in light field microscopy. The results and approach we have presented here are potentially applicable to any multi-aperture computational imaging system with a space-variant PSF endowed with translational symmetry properties. We have made available an open-source implementation of our algorithms (9), with performance measured to be more than an order of magnitude faster than previously-available codes. We anticipate that our algorithms and opensource implementation will lead to an expansion in the uptake of light field microscopy as a tool for 3D+time imaging of rapid biological processes, now that the excessive computational demands of the reconstruction process have been brought under control.

## Data availability

The computer code and test datasets underlying the results presented in this paper are available at Refs. (9, 12).

